# NPP-21/TPR is required for developmental control of spindle checkpoint strength in *C. elegans*

**DOI:** 10.64898/2026.04.13.718277

**Authors:** Natalie Gallagher, Shevelle Brown, Valentin Duprat, Simone Köhler, Abby F. Dernburg, Needhi Bhalla

## Abstract

The spindle checkpoint ensures accurate chromosome segregation by monitoring whether chromosomes, via kinetochores, are properly attached to the spindle. If chromosomes fail to establish bipolar attachment, the checkpoint delays the cell cycle to enable error correction. In *C. elegans* early embryos, activation of the spindle checkpoint produces a longer mitotic delay in primordial germ cells than somatic cells. We show that the conserved nucleoporin and spindle matrix component, NPP-21/TPR, is required for the stronger spindle checkpoint in germline cells. A checkpoint-proficient NPP-21::GFP transgene localizes to a spindle-like structure during mitosis and is enriched in germline cells, consistent with a cell-fate specific function for this protein. Finally, NPP-21 controls spindle checkpoint strength in germline cells via two, potentially linked, mechanisms: concentrating PCH-2 around mitotic chromosomes and promoting the localization of the checkpoint effector, Mad2, to unattached kinetochores. These experiments demonstrate a developmental role for NPP-21, and the spindle matrix, in controlling spindle checkpoint strength in immortal germline cells in *C. elegans*.

## Introduction

The spindle checkpoint is a cell cycle surveillance mechanism that ensures that chromosomes are attached to the mitotic or meiotic spindle to guarantee accurate chromosome segregation and prevent aneuploidy. When chromosomes are unattached, a soluble, molecular signal halts or delays the cell cycle to allow for the defect to be corrected. Defects in spindle checkpoint components are associated with cancer progression (Kops et al., 2005; Matthews et al., 2022) and infertility (Baker et al., 2004; Huang et al., 2023; Zhang et al., 2020), highlighting their importance to human health.

The spindle checkpoint initiates from kinetochores, the specialized structures that assemble at centromeres to mediate microtubule attachment and proper chromosome segregation (Lara-Gonzalez et al., 2021). A complex of the conserved checkpoint factors, Mad1 and Mad2, binds unattached or incorrectly attached kinetochores. Mad2 adopts two functionally distinct conformations: an open, inactive form and a closed, active form. When Mad2 binds a specific motif, called the closure motif, in its binding partners, such as Mad1 and Cdc20, Mad2 undergoes a conformational conversion from inactive (open) to active (closed). At unattached kinetochores, the Mad1/Mad2 complex recruits additional inactive Mad2 so that it can bind Cdc20, adopting the active conformation and forming the Mitotic Checkpoint Complex, or MCC. The MCC sequesters Cdc20, a factor required to promote anaphase, to inhibit the cell cycle and allow time for error correction (Lara-Gonzalez et al., 2021). The evolutionarily ancient ATP-ase PCH-2/TRIP13 promotes spindle checkpoint function by ensuring the availability of open, inactive Mad2 (Ma and Poon, 2016; Ma and Poon, 2018; Nelson et al., 2015)

Despite the important role of the spindle checkpoint in preventing aneuploidy, the mitotic delay or arrest installed by the spindle checkpoint can be highly variable. The variability of the cell cycle delay is called spindle checkpoint strength and depends on the number of unattached kinetochores, cell volume, and cell fate (Collin et al., 2013; Galli and Morgan, 2016; Gerhold et al., 2018; Kyogoku and Kitajima, 2017). For example, during early embryogenesis in *C. elegans*, primordial germ cells have a stronger checkpoint than their somatic counterparts, independent of their smaller size (Galli and Morgan, 2016; Gerhold et al., 2018), demonstrating developmental differences in checkpoint strength. We showed that the stronger checkpoint in germline cells depends on the asymmetric enrichment of the spindle checkpoint factor, PCH-2/TRIP13 (Defachelles et al., 2020).

During mitosis in early *C. elegans* embryos, following nuclear envelope breakdown, PCH-2::GFP and other checkpoint components, such as GFP::Mad1 and GFP::Mad2, remain localized around mitotic chromosomes (Defachelles et al., 2020; Essex et al., 2009). This diffuse localization resembles a structure known as the spindle matrix (Zheng, 2010). While the composition and function of the spindle matrix remain controversial, some characterized spindle matrix components interact with spindle checkpoint components (Lince-Faria et al., 2009; Schweizer et al., 2013), making them attractive candidates to regulate spindle checkpoint strength during development.

NPP-21/TPR is the *C. elegans* ortholog of the *Drosophila* spindle matrix protein, Megator, and the human nucleoporin TPR. In *C. elegans*, NPP-21/TPR has been primarily characterized as a nucleoporin, a member of a large family of proteins that form the nuclear pore and allow for macromolecular transport between the cytoplasm and nucleus (Cohen-Fix and Askjaer, 2017). NPP-21/TPR forms the nucleoplasmic face or basket of the pore (Cohen-Fix and Askjaer, 2017). In *Drosophila*, Megator is a nucleoporin that also localizes to the area occupied by the miotic spindle (Qi et al., 2004) and supports robust recruitment of Mad2 to unattached kinetochores (Lince-Faria et al., 2009).

Here, we show that NPP-21/TPR, in its capacity as a spindle matrix component, is required for developmental control of spindle checkpoint strength in *C. elegans*. A strain carrying both a transgene of *npp-21*, *AID-3xFLAG::npp-21*, and the Arabidopsis TIR1 gene is functional for checkpoint activation in somatic cells of the early embryo but cannot support the stronger checkpoint in germline cells. When NPP-21 is acutely degraded, checkpoint strength is further compromised in germline cells but has no effect on somatic cells. Consistent with this cell-fate specific function, NPP-21::GFP consistently forms a structure that resembles the spindle matrix during mitosis and is asymmetrically enriched in germline cells, compared to somatic cells.

Finally, NPP-21 ensures a stronger checkpoint in germline cells by concentrating PCH-2::GFP around mitotic chromosomes and promoting the robust recruitment of GFP::Mad2 to unattached kinetochores. We propose that during the rapid cell divisions of early embryogenesis, NPP-21, and the spindle matrix, concentrate checkpoint factors around mitotic chromosomes to ensure genomic integrity, a role that is particularly important in primordial germ cells that will give rise to sperm and eggs.

## Results and Discussion

### NPP-21 is required for the stronger spindle checkpoint in germline, P_1_ cells

NPP-21 is an essential gene (Piano et al., 2000). Therefore, to determine whether NPP-21 plays a role in the spindle checkpoint in early developing embryos, we exploited the auxin-inducible degron system (AID) to acutely degrade NPP-21 in oocytes and early embryos. Using CRISPR/Cas9, we generated a version of NPP-21 tagged at its N-terminus with the AID degron and a 3xFLAG tag (*AID-3xFLAG::npp-21*, also referred to as *AID::npp-21*) in a strain that also expresses the Arabidopsis TIR1 protein via a germline promoter (*Pgld-1::TIR1-mRuby)* (Zhang et al., 2015). In the presence of auxin, TIR1 binds to degron-tagged proteins and targets them for polyubiquitylation and degradation by the proteasome (Nishimura et al., 2009). In the presence of ethanol and absence of auxin, this strain produces modest defects in in meiotic chromosome segregation, illustrated by the elevated frequencies of inviable embryos and male self-progeny (Table 1). When these worms were exposed to auxin, they produced few viable progeny and 25% male self-progeny (Table 1). Thus, the *AID::npp-21* transgene is mostly functional, with some slight defects in embryonic viability and meiotic chromosome segregation, which may be the product of the tag and/or auxin-independent degradation. By contrast, auxin-induced depletion of AID-3xFLAG::NPP-21 results in very high embryonic lethality and severe defects in meiotic chromosome segregation, consistent with its essential role (Piano et al., 2000).

**Table 1:**
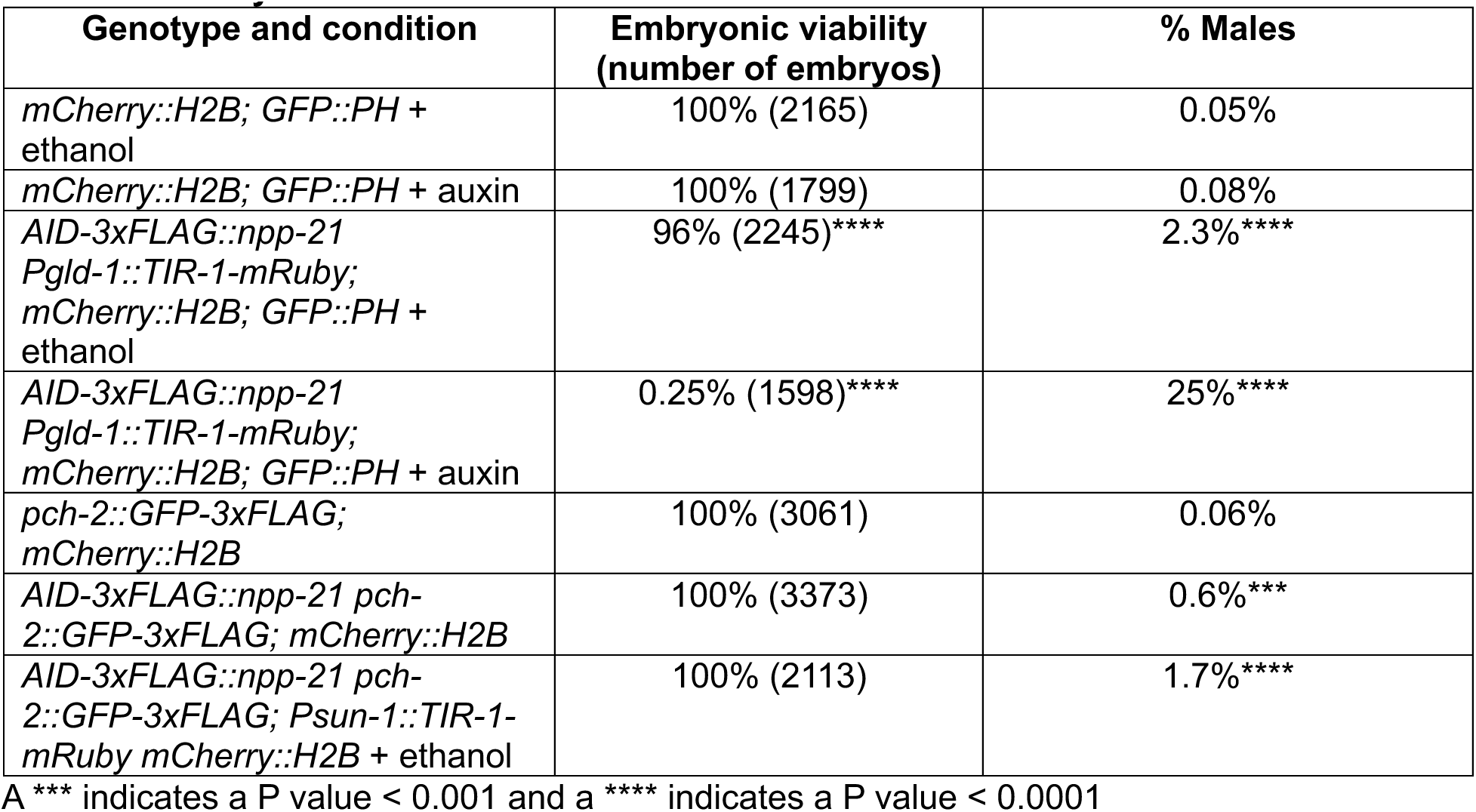
Viability of strains.

We tested whether we could rapidly degrade AID-3xFLAG::NPP-21. We performed this experiment in the most proximal oocytes undergoing cellularization (-1 oocyte). Upon fertilization, these oocytes complete meiosis and enter mitosis to begin embryonic development within an hour (Robertson and Lin, 2013). We exposed young adult hermaphrodites worms undergoing oogenesis to auxin for one hour and performed indirect immunofluorescence against the FLAG tag to visualize AID-3xFLAG::NPP-21. In worms exposed to ethanol, we observed staining at the nuclear periphery in the most proximal oocytes, consistent with its known role as a member of the nuclear pore complex (Figure 1A). After one hour of auxin treatment, we could no longer detect AID-3xFLAG::NPP-21 at the nuclear periphery (Figure 1A), indicating that we can reliably and rapidly degrade NPP-21 in oocytes that are about to be fertilized and develop into embryos.

**Figure 1:**
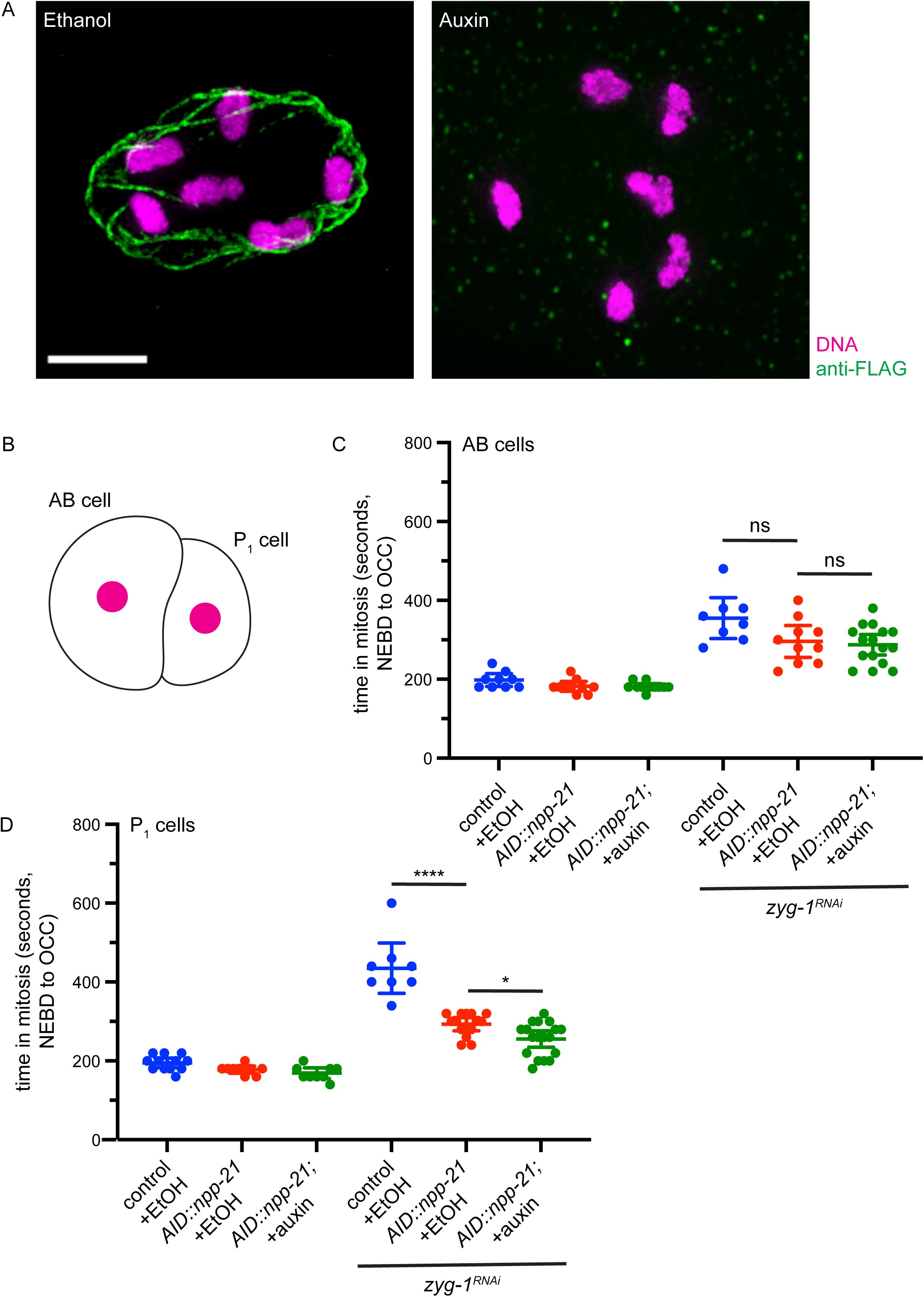
NPP-21 is required for the stronger spindle checkpoint in germline cells of the two-cell embryo. A. AID-3xFLAG::NPP-21 is degraded in oocytes after one hour of auxin treatment. Images of cellularized (-1) oocytes from worms treated with ethanol or auxin stained with antibodies against the FLAG tag and DAPI to visualize DNA. Scale bar indicates 5 microns. B. Cartoon indicating a two cell embryo with the AB and P_1_ cell labeled and the nucleus colored magenta. C. AID-3XFLAG::NPP-21 is functional and not required for the spindle checkpoint in AB, somatic, cells. Graph depicting mitotic timing in AB cells of embryos treated with ethanol or auxin and control RNAi or *zyg-1* RNAi. D. AID-3XFLAG::NPP-21 cannot support the stronger checkpoint and is required for a robust spindle checkpoint in P_1_, germline, cells. Graph depicting mitotic timing in P_1_ cells of embryos treated with ethanol or auxin and control RNAi or *zyg-1* RNAi. In all graphs, a * indicates a p value < 0.05, a ** indicates a p value < 0.01, a *** indicates a p value < 0.001, a **** indicates a p value < 0.0001, ns indicates not significant and error bars indicate 95% confidence intervals.

Having established this protocol, we used it to test whether degradation of NPP-21 affected normal mitotic timing and spindle checkpoint function in early embryos, specifically two cell embryos which have already differentiated into soma (AB cells) and germline (P_1_ cells) (Figure 1B). We analyze mitotic timing by calculating the time between nuclear envelope breakdown (NEBD) and the onset of cortical contractility (OCC), which coincides with anaphase in embryos expressing an mCherry tagged histone, mCherry::H2B, and the plasma membrane marker, GFP::PH (Essex et al., 2009). NEBD is visualized as the diffusion of soluble mCherry::H2B signal from the nucleus and OCC is detected when the plasma membrane changes conformation from circular to rectangular (Supplemental Figure 1A). In P_1_ cells, it is difficult to assess OCC and we used anaphase as an indicator of mitotic exit (Supplemental Figure 1A).

When transgenic *AID::npp-21* worms were exposed to ethanol or auxin, we did not detect any difference in mitotic timing between ethanol-treated or auxin-treated worms, or with control worms treated with ethanol, in either AB or P_1_ cells, indicating that NPP-21 is not required for unperturbed mitosis in either cell type (Figures 1C and D).

Next, we activated the spindle checkpoint in *AID::npp-21* AB cells by performing RNA interference (RNAi) against *zyg-1*, a conserved cell cycle kinase required for centrosome duplication (O’Connell et al., 2001). In the absence of *zyg-1*, two cell embryos only have one centrosome and generate a monopolar spindle, producing chromosomes with unattached kinetochores (Essex et al., 2009). In *zyg-1^RNAi^* embryos, OCC is defined as the formation of persistent membrane blebs between the AB and P_1_ cells (arrows in Supplemental Figure 1A). In AB cells of *zyg-1^RNAi^* embryos, we observed a slight decrease in mitotic timing in transgenic *AID::npp-21* worms on ethanol or auxin (Figure 1C), when compared to control embryos on ethanol. This difference was not statistically significant, indicating that NPP-21 might play a role in promoting a strong spindle checkpoint in AB cells but is not required for the checkpoint in these cells. In contrast, when we activated the spindle checkpoint in P_1_ cells in *AID::npp-21* worms exposed to ethanol, these cells showed similar mitotic timing as AB cells treated with *zyg-1* RNAi (Figures 1C and D) and a weaker checkpoint than control worms (Figure 1D).

These data indicate that either the transgenic *AID::npp-21* is not fully functional for spindle checkpoint function in P_1_ cells or that the presence of TIR1, even in the absence of auxin, affects protein function. More importantly, these data suggest a role for NPP-21 in spindle checkpoint strength in this specific cell type. When *AID::npp-21* worms are exposed to auxin for more than one hour, we detected a further weakening of the spindle checkpoint (Figure 1D), indicating that NPP-21 is required for a robust spindle checkpoint specifically in P_1_ cells.

### NPP-21::GFP adopts a structure similar to the spindle matrix and is enriched in P_1_ cells

Given this cell fate-specific difference in the requirement for NPP-21 function in the spindle checkpoint, we localized NPP-21::GFP in AB and P_1_ cells in two cell embryos (Thomas et al., 2023). First, we verified that *npp-21::gfp* worms were competent for checkpoint activation in AB and P_1_ cells. Because the presence of NPP-21::GFP prevented us from using GFP::PH to monitor OCC, we measured mitotic timing using mCherry::H2B from NEBD to decondensation of mitotic chromosomes (DECON) (Supplemental Figure 1B) (Essex et al., 2009). In embryos expressing NPP-21::GFP, both AB and P_1_ cells activated the spindle checkpoint and P_1_ cells displayed a stronger checkpoint than AB cells, similar to control worms (Figure 2A), verifying that NPP-21::GFP is fully functional for checkpoint activation. Next, we localized NPP-21::GFP and observed a striking difference in its localization in AB and P_1_ cells. In 75% of two-cell embryos (15/20), NPP-21::GFP remained in a cloud around mitotic chromosomes 120 seconds after NEBD, a timepoint that corresponds to metaphase, in AB cells (see Materials and Methods). In contrast to AB cells, after NEBD in P_1_ cells, NPP-21::GFP reorganized around mitotic chromosomes in a structure that resembled the mitotic spindle, similar to a spindle matrix. In 25% of two cell embryos, both AB and P_1_ cells exhibited this spindle-like localization (Figure 2B). Thus, in P_1_ cells 100% of the time, and in AB cells 25% of the time, NPP-21::GFP adopts a localization pattern during mitosis that resembles the spindle matrix (see Supplemental Movies).

**Figure 2:**
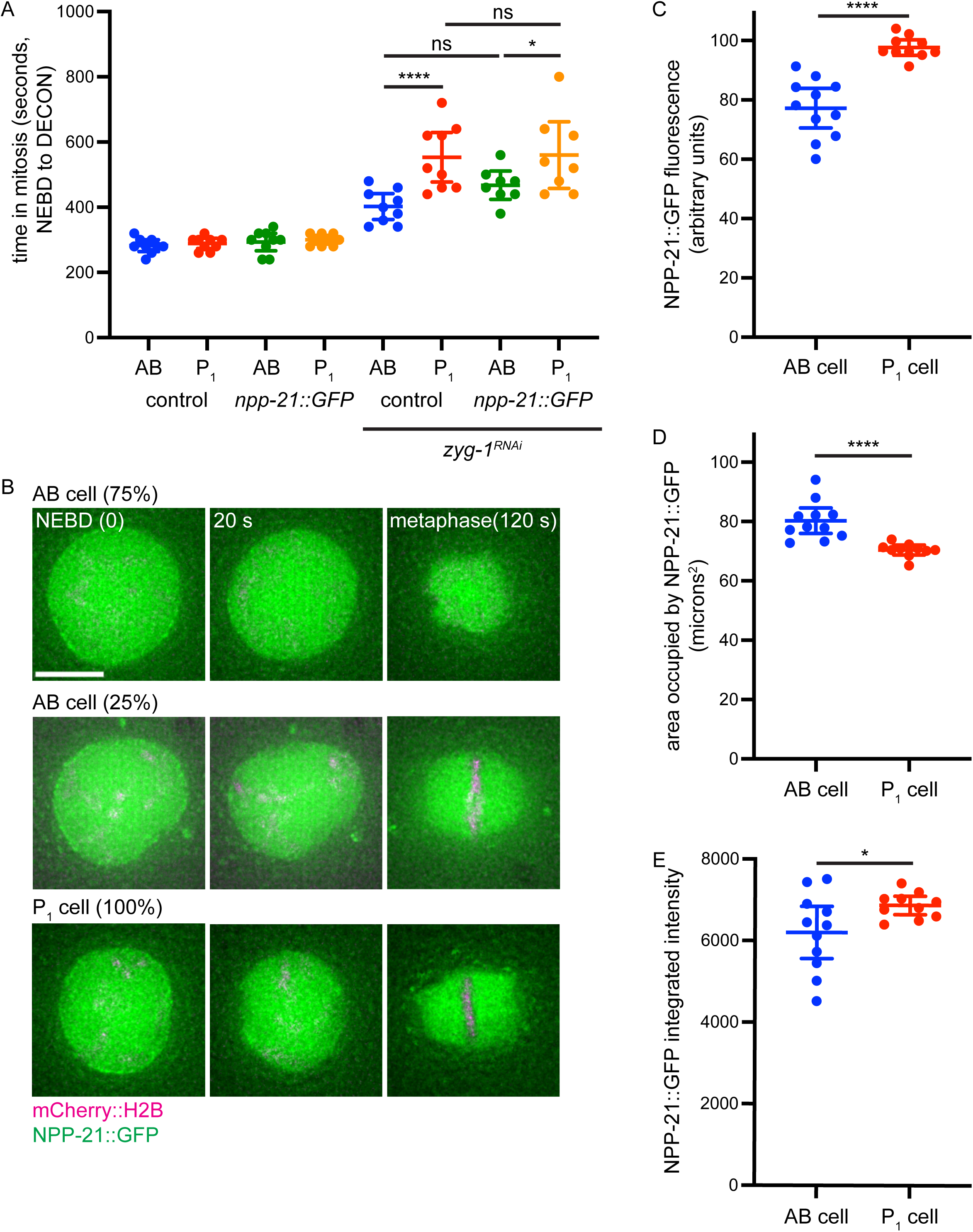
NPP-21::GFP is enriched in P_1_, germline, cells and its localization is distinct from that in AB, somatic, cells. A. NPP-21::GFP is functional for the spindle checkpoint in AB and P_1_ cells. A graph depicting mitotic timing in AB and P_1_ cells of control and *npp-21::GFP* animals treated with control and *zyg-1* RNAi. B. NPP-21::GFP localization in P_1_ cells is different than in AB cells in most two cell embryos. Images of AB and P_1_ cells expressing mCherry::H2B (magenta) and NPP-21::GFP (green) at NEBD, 20 seconds after NEBD and at metaphase (120 seconds after NEBD). C. NPP-21::GFP is asymmetrically enriched in P_1_ cells. Graph depicting average fluorescence of NPP-21::GFP in AB and P_1_ cells 20 seconds after NEBD. D. The area occupied by NPP-21::GFP is smaller in P_1_ cells. Graph depicting the area in microns^2^ of NPP-21::GFP in AB and P_1_ cells 20 seconds after NEBD. E. NPP-21::GFP is asymmetrically enriched in P_1_ cells. Graph depicting integrated intensity of NPP-21-::GFP in AB and P_1_ cells 20 seconds after NEBD.

We previously showed that PCH-2::GFP, which is required for spindle checkpoint function, is enriched in P_1_ cells (Defachelles et al., 2020). To determine whether NPP-21::GFP is also enriched in P_1_ cells, we quantified NPP-21::GFP fluorescence immediately following NEBD (images labeled 20 seconds in Figure 2B). We performed this quantification at this stage of the cell cycle since NPP-21::GFP localization was similar in both AB and P_1_ cells immediately after NEBD and entry into mitosis, in contrast to the changes in localization we observed at metaphase in AB and P_1_ cells (Figure 2B). When we performed this analysis, we detected more NPP-21::GFP in P_1_ cells than AB cells (Figure 2C). We were concerned that, even at this early timepoint, the area occupied by NPP-21::GFP was different in AB and P_1_ cells, potentially affecting this quantification. To test this, we determined the area occupied by NPP-21::GFP signal in AB and P_1_ cells and found that this area was significantly smaller in P_1_ cells than AB cells (Figure 2D). Since AB and P1 cells show similar nuclear areas prior to NEBD (Gerhold et al., 2018), this indicates that NPP-21::GFP is regulated differently in P_1_ cells even at this early timepoint. However, when we calculated the integrated intensity of NPP-21::GFP signal (Figure 2E) in both cell types, NPP-21::GFP signal was still significantly higher in P_1_ cells than AB cells, indicating that NPP-21::GFP is enriched in P_1_ cells, potentially to drive the stronger checkpoint.

### NPP-21 concentrates PCH-2::GFP around mitotic chromosomes

PCH-2 and its mammalian ortholog, Trip13, remodels active Mad2 to generate inactive Mad2 (Ye et al., 2015). This remodeling guarantees the availability of inactive Mad2 during checkpoint activation so that the conversion of inactive Mad2 to active Mad2, and the production of soluble MCC, is the rate limiting event in spindle checkpoint activation (Ma and Poon, 2016; Ma and Poon, 2018; Nelson et al., 2015).

To evaluate whether NPP-21 contributed to the stronger spindle checkpoint in P_1_ cells by regulating PCH-2, we generated a strain that expresses both AID::NPP-21 and a fluorescently tagged, functional PCH-2::GFP-3xFLAG (Nelson et al., 2015). Both *npp-21* and *pch-2* loci are closely linked (0.20 cM apart). To generate the *AID::npp-21 pch-2::GFP-3xFLAG* double mutant strain, the TIR1 transgene, also on the same chromosome, was crossed off in the recombination event. Since we observed the defect in checkpoint strength in P_1_ cells even in embryos that have not been treated with auxin (Figure 1D), we first determined whether we could perform this experiment in worms that did not express the TIR1 transgene. These double mutants do not exhibit any embryonic inviability, indicating that the embryonic inviability we observe in *AID-3xFLAG::npp-21 Pgld-1::TIR1-mRuby* strains on ethanol is likely the product of auxin-independent but TIR1-dependent degradation of NPP-21 (Table 1). *AID-3xFLAG::npp-21 pch-2::GFP-3xFLAG* hermaphrodites produced males at a higher, statistically significant frequency than *pch-2::GFP-3xFLAG* animals, suggesting a very weak defect in meiotic chromosome segregation (Table 1). However, *AID-3xFLAG::npp-21 pch-2::GFP-3xFLAG* embryos display a stronger checkpoint in P_1_ cells, similar to control animals (Figure 3A), indicating that *AID-3xFlag::npp-21* is fully functional and the weaker checkpoint we observed in *3xFLAG::npp-21 Pgld-1::TIR1-mRuby* P1 cells was a product of TIR1 expression.

**Figure 3:**
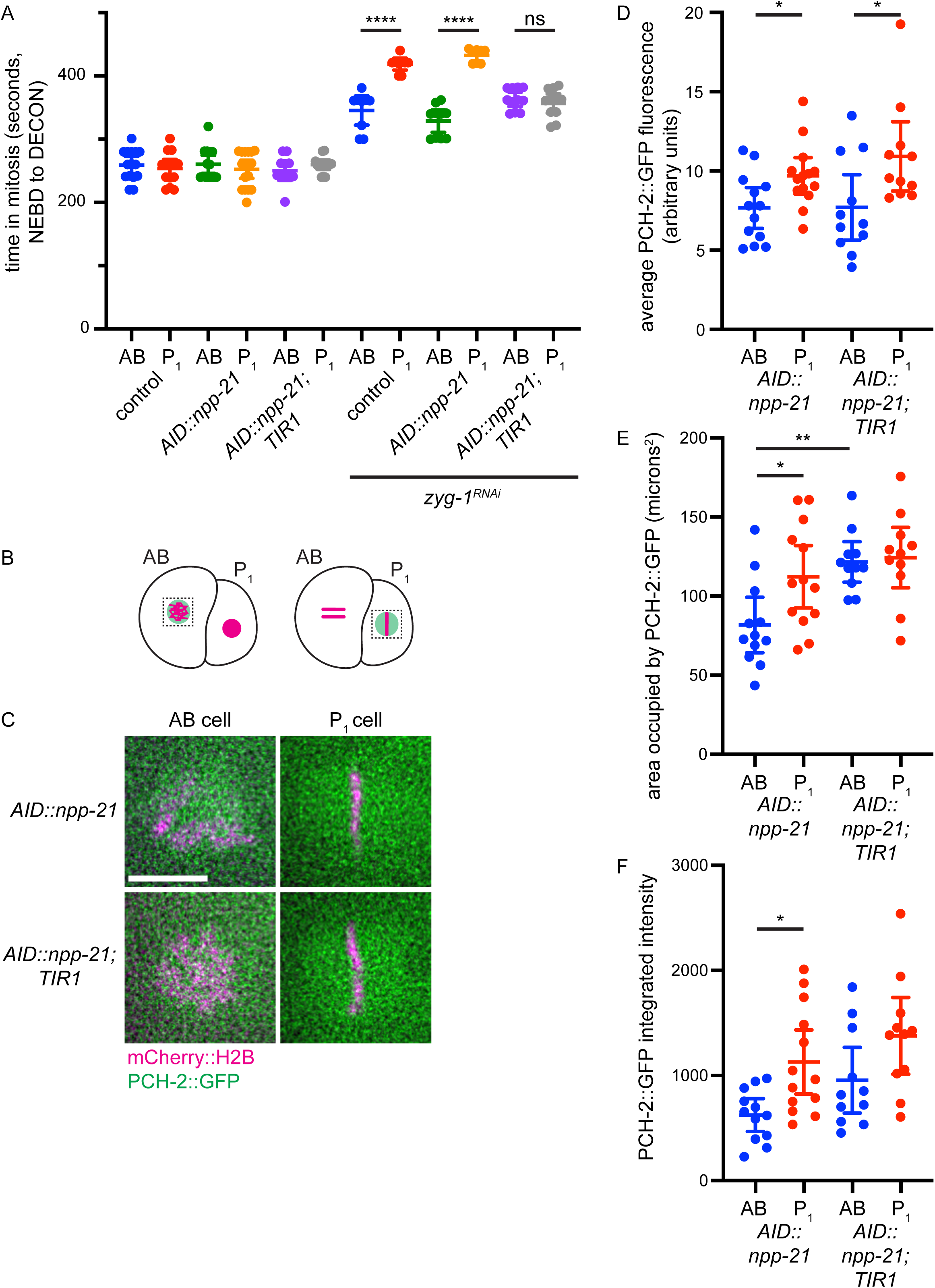
NPP-21 concentrates PCH-2::GFP around mitotic chromosomes in both AB and P_1_ cells. A. Worms expressing both AID-3XFLAG::NPP-21 and TIR1 cannot support the stronger checkpoint. A graph depicting mitotic timing in AB and P_1_ cells of control, *AID::npp-21* and *AID::npp-21;TIR1* animals treated with control and *zyg-1* RNAi. B. Cartoon of PCH-2::GFP localization in AB and P_1_ cells. C. Images of PCH-2::GFP (green) localization in AB and P_1_ cells expressing mCherry::H2B (magenta) in metaphase, in embryos without and with the TIR1 gene. Scale bar indicates 5 microns. D. PCH-2::GFP is asymmetrically enriched in P_1_ cells, in embryos without and with the TIR1 gene. Graph depicting average fluorescence of PCH-2::GFP in AB and P_1_ cells at metaphase. E. The area occupied by PCH-2::GFP is greater in AB and P_1_ cells in embryos expressing TIR1. Graph depicting the area in microns^2^ of PCH-2::GFP in AB and P_1_ cells in metaphase. E. PCH-2::GFP is less concentrated in AB and P_1_ cells of embryos expressing TIR1. Graph depicting integrated intensity of PCH-2::GFP in AB and P_1_ cells.

We introduced a different TIR1 transgene controlled by another germline promoter, *Psun-1::TIR1-mRuby,* into strains with both *AID-3xFLAG::npp-21* and *pch-2::GFP-3xFLAG* transgenes. Strains carrying AID::NPP-21, this transgenic version of TIR1 and PCH-2::GFP showed a weaker spindle checkpoint in P_1_ cells, similar to AB cells and in contrast to control animals and strains with only AID::NPP-21 and PCH-2::GFP (Figure 3A). This phenotype was also similar to the original strain expressing AID::NPP-21 and TIR1 via the *gld-1* promoter (Figure 1D). When these worms were grown on plates containing ethanol, we observed an increase in male self-progeny and no effect on viability (Table 1).

We quantified PCH-2::GFP in both AB and P_1_ cells in embryos expressing AID::NPP-21 and PCH-2::GFP with or without TIR1 (Figures 3B and C). Both strains showed statistically significant enrichment of PCH-2::GFP in P_1_ cells, compared to AB cells (Figure 3D). However, we noticed that the area occupied by PCH-2::GFP increased in both AB and P_1_ cells in strains expressing TIR1, compared to strains without TIR1 (Figure 3E and Supplemental Figure 2).

Moreover, in P_1_ cells without TIR1, the area occupied by PCH-2::GFP was larger than that observed in AB cells without TIR1, likely because PCH-2::GFP often adopted a matrix-like structure in P_1_ cells (55%, 6 of 11 embryos) (Figure 3C and Supplemental Figure 2). However, this structure was much less defined than the one observed with NPP-21::GFP (Figure 2B).

When the area occupied by PCH-2::GFP was used to calculate the integrated intensity of PCH-2::GFP in AID::NPP-21 embryos with and without TIR1, P_1_ cells in strains with TIR1 still had higher levels of PCH-2::GFP but the comparison with AB cells was not statistically significant (Figure 3F). This was in direct contrast to the experiments performed in embryos without TIR1 (Figure 3F). These data indicate that NPP-21 is not required to enrich PCH-2::GFP in P_1_ cells but instead, promotes its concentration around mitotic chromosomes in both AB and P_1_ cells.

Given the defect in checkpoint strength in P_1_ cells in this strain, we would argue that this concentration is particularly important for the stronger checkpoint in P_1_ cells. The inability to properly concentrate PCH-2::GFP in P_1_ cells may also explain why PCH-2::GFP rarely adopts a matrix-like structure in P_1_ cells in embryos expressing AID::NPP-21 and TIR1 (18%, 2 of 11 embryos) (Figure 3C and Supplemental Figure 2).

### NPP-21 promotes GFP::MAD-2 recruitment to unattached kinetochores in P_1_ cells

In both cultured *Drosophila* and human (HeLa) cells, NPP-21 orthologs, Megator and TPR respectively, promote the localization of Mad2, but not Mad1, to unattached kinetochores by acting as spatial regulators or ensuring protein stability (Lince-Faria et al., 2009; Schweizer et al., 2013). To determine if NPP-21 plays a similar role in *C. elegans* embryos (Figures 4A and B), we quantified GFP::MAD-2 at unattached kinetochores in *AID::npp-21* embryos (Figures 4B and C). In AB cells, GFP::MAD-2 recruitment to unattached kinetochores was unaffected by NPP-21 depletion (Figures 4B and C), consistent with their normal checkpoint activation (Figure 1C). P_1_ cells treated with ethanol also showed normal GFP::MAD-2 recruitment, indicating that the weaker spindle checkpoint in P_1_ cells is not the result of a baseline defect in localizing Mad2 (Figures 4B and C). By contrast, NPP-21 depletion sharply reduced GFP::MAD-2 localization to unattached kinetochores in P_1_ cells (Figures 4B and C), demonstrating that the weaker checkpoint response in these cells stems from the inability to robustly localize Mad2 to unattached kinetochores.

**Figure 4:**
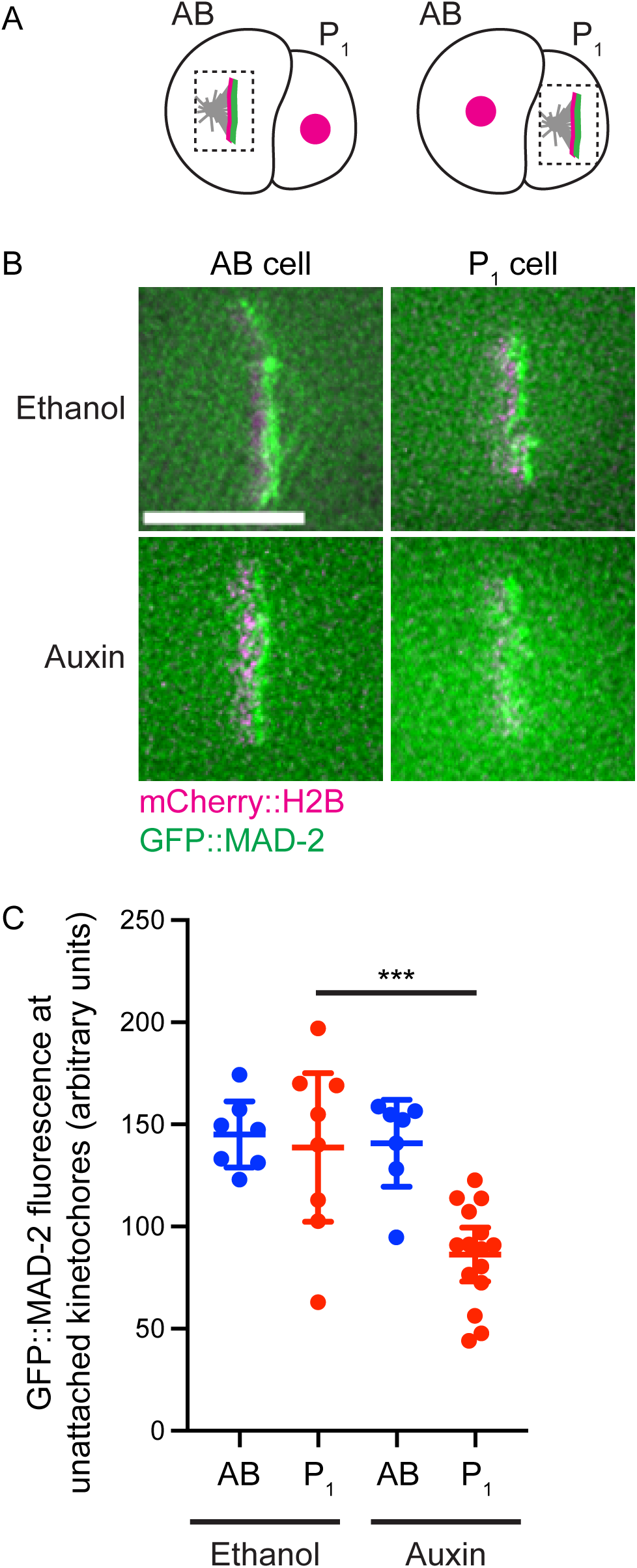
NPP-21 is required for GFP::MAD-2 recruitment to unattached kinetochores in P_1_, germline, cells. A. Cartoon of GFP::MAD-2 localization in AB and P_1_ cells in *zyg-1^RNAi^* embryos. B. Images of GFP::MAD-2 (green) localization in AB and P_1_ cells in *zyg-1^RNAi^* embryos expressing mCherry::H2B (magenta), treated with ethanol (top) or auxin (bottom). Scale bar indicates 5 microns. C. Quantification of GFP::MAD-2 in AB and P_1_ cells at unattached kinetochores in *zyg-1^RNAi^*embryos treated with ethanol or auxin.

During early development, embryos balance the rapid cell divisions of early embryogenesis with the imperative of maintaining genomic integrity (Fusari and Milan, 2026), particularly in immortal germline cells that will give rise to sperm and eggs. NPP-21, through its spindle matrix function, contributes to this balance by promoting a stronger spindle checkpoint response in germline cells, in contrast to somatic cells. The NPP-21 protein displays germline specific behavior: it is enriched and consistently forms a spindle matrix in P_1_ cells, suggesting a conserved developmental role during embryogenesis. This function acts at two control points: concentrating PCH-2 around mitotic chromosomes and promoting Mad2 recruitment to unattached kinetochores. Because PCH-2 promotes the formation of inactive, open Mad2 for checkpoint signaling, these two control points may be mechanistically linked. Whether other, conserved spindle matrix components play similar, developmental, roles during embryogenesis is an interesting, open question for our future studies.

## Materials and Methods

### *C. elegans* Strains and Husbandry

The wildtype *C. elegans* strain background was Bristol N2 (Brenner, 1974) . All strains were maintained at 20**°**C. See Supplemental Table 1 for a list of all *C. elegans* strains used in this study.

The *AID::3xFLAG::npp-21* transgenic strain was generated by CRISPR/Cas9 genomic editing (Paix et al., 2017). Young adult worms carrying the *Pgld-1::Tir1mRuby::3’UTR(gld-1) (ieSi64)* transgene were injected with preassembled Cas9-RNP, 23 ng/μL of ssDNA repair template and 5 ng/uL pCFJ90 and 2.5 ng/uL pCFJ104 as co-injection markers. The target crRNA was TAGTCTTTCAGTTATGGATG (IDT) and the ssDNA repair template was generated by primer extension PCR using the following primer GTGTTATCTAATTAAATCTCTCAGTTCCTTCAAGTTATCATAT and gBlock CTCTCAGTTCCTTCAAGTTATCATATTTTTTAAGTAAAACTAATCGCCCGAAATTTAGTCTTTCAGTTATGCCTAAAGATCCAGCCAAACCTCCGGCCAAGGCACAAGTTGTGGGATGGCCACCGGTGAGATCATACCGGAAGAACGTGATGGTTTCCTGCCAAAAATCAAGCGGTGGCCCGGAGGCGGCGGCGTTCGTGAAGGACTATAAAGATCACGACGGAGATTACAAGGACCATGATATCGACTACAAGGACGACGACGACAAGGGAGATGTGGATGCGCCACTTCAGGCGCCTGAACAACCGGTCGCAGACGCTGACGATGAAGAAGCTAACTGGGAAATGGAAAAAGCCGAAATGAAACGGATTGAG (both ordered from IDT). Transformants were identified and worms carrying the transgene were genotyped using the following primers: AATTGGCATCGCTGATCAATGAG and GCTCTTTAATTTCAGCCGCATGAC. The presence of the insertion was verified by sequencing and the transgenic strain was backcrossed at least three time before analysis.

### Viability and fertility

To score total progeny and male self-progeny, L4 hermaphrodites were picked onto individual plates with 1mM auxin, ethanol or no additional treatment, and transferred to new plates daily over 4 days. The eggs laid on each plate were counted after removing the parent. Viable progeny and male progeny were quantified when the F1 reached L4 or adult stages (2-3 days post egg laying).

### Microscopy & Mitotic Timing Experiments

All immunofluorescence and live microscopy was performed on a DeltaVision Personal DV deconvolution microscope (Applied Precision) equipped with a 100X N.A. 1.40 oil-immersion objective (Olympus) coupled with a CoolSNAP charge-coupled camera (Roper Scientific). For live microscopy of two cell embryos, eggs were dissected 18-26 hours post-L4 into 1X Egg Buffer (25mM Hepes pH7.4, 118uM NaCl, 48mM KCl, 2mM EDTA, 0.5mM EGTA) and mounted on 2% agarose pads for immediate analysis. Environmental temperature was approximately 21**°**C during image collection. For mitotic timing experiments, Z-sections were acquired with 8x 2μm steps using a 100X objective (Olympus) at 20s intervals. Exposure time was 100ms for mCherry::H2B and 50ms for GFP::PH. Mitotic duration was calculated for the AB cell in the presence of monopolar spindles as the interval between NEBD to the onset of cortical contractility (OCC) or the interval between NEBD and chromosome decondensation (DECON). NEBD was defined by the equilibration of mCherry::H2B from the nucleus into the cytosol. OCC was defined as the change in conformation of the plasma membrane from circular to rectangular in AB cells, as anaphase in P_1_ cells or as the first frame when a persistent membrane bleb formed from the cortex of the embryo in *zyg-1^RNAi^*embryos (Supplemental Figure 1A). DECON was defined as the loss of punctate mCherry::H2B signal within the decondensing chromatin (Supplemental Figure 1B). To minimize bleaching and maximize signal intensity of GFP-tagged spindle checkpoint components (NPP-21, PCH-2 and MAD-2), imaging was started just after NEBD as visualized by mCherry::H2B. Here, 8x 1μm steps were captured with 250ms GFP and 100ms mCherry exposures at 20s intervals.

Immunostaining was performed on worms 20–24 h after L4 stage, as described in (Russo et al., 2021). Gonad dissections were performed in 1× EBT (250 mM Hepes-Cl, pH 7.4, 1.18 M NaCl, 480 mM KCl, 20 mM EDTA, and 5 mM EGTA) + 0.1% Tween 20 and 20 mM sodium azide. An equal volume of 7.4% formaldehyde in EBT (final concentration was 3.7% formaldehyde) was added and allowed to incubate under a coverslip for 5 min. The sample was mounted on HistoBond slides (75 × 25 × 1 mm from VWR), freeze-cracked, and incubated in methanol at −20°C for slightly more than 1 min and transferred to PBST (PBS with Tween 20). After several washes of PBST, the samples were incubated for 30 min in 1% bovine serum albumin diluted in PBST. A hand-cut paraffin square was used to cover the tissue with 50 µl of antibody solution. Incubation was conducted in a humid chamber overnight at 4°C. Slides were rinsed in PBST and then incubated for 2 h at room temperature with fluorophore-conjugated secondary antibody at a dilution of 1:500. Samples were rinsed several times and DAPI stained in PBST, then mounted in 13 µl of mounting media (20 M *N*-propyl gallate [Sigma-Aldrich] and 0.14 M Tris in glycerol) with a no. 1 1/2 (22 mm^2^) coverslip, and sealed with nail polish.

Mouse anti-FLAG [Sigma] primary antibodies were used for immunofluorescence at 1:1000 and Alexa-Fluor 488 anti-mouse (Invitrogen) secondary antibodies were used at 1:500. Three-dimensional image stacks were collected at 0.2-μm Z-spacing and processed by constrained, iterative deconvolution. Image scaling and analysis were performed using functions in the softWoRx software package. Projections were calculated by a maximum intensity algorithm. Composite images were assembled and some false coloring was performed with Adobe Photoshop.

### Quantification of NPP-21::GFP, GFP::MAD-2, and PCH-2::GFP-3XFLAG

Analysis was performed in Fiji. Quantification of fluorescence around mitotic chromosomes was quantified as described in (Defachelles et al., 2020). Sum intensity projections were generated and average fluorescence in the area around mitotic chromosomes was measured in Fiji. Background average fluorescence was measured in a 30-pixel band around this “cloud” and subtracted from the initial fluorescence intensity to determine the final value. In some of our movies to quantify PCH-2::GFP and localize NPP-21::GFP, identifying a clear metaphase plate was more difficult in AB than in P1 cells. Therefore, to ensure that we were performing these experiments at the same stage in mitosis in these two cell types, frames for the relevant analysis were normalized relative to NEBD and mitotic exit.

Quantification of unattached kinetochore signal was performed essentially as described for GFP::MAD-2 quantification in (Nelson et al., 2015). Maximum intensity projections of both mCherry::H2B and GFP fusion proteins were made after the pseudo-metaphase plate was generated. The image was rotated so the metaphase plate was vertical, channels were split, and the maximum GFP pixel was identified using the process function within a box on the unattached side of the metaphase plate. In the same x-plane, the maximum mCherry::H2B pixel was found. The width was changed to 12 pixels and the maximum GFP signal intensity was recorded in this 12 pixel window centered at the mCherry maxima. The background GFP signal was calculated by taking the average GFP intensity of a 4 pixel box in the same x-plane, 8 pixels away from the maximum mCherry on the opposite side of the pseudo-metaphase plate to the maximum GFP (i.e. the attached side). This background GFP was then subtracted from the maximum to measure the kinetochore bound GFP fusion intensity. This process was repeated at least 7x for each genetic background and treatment.

### Feeding RNA interference (RNAi)

RNA interference (RNAi) was performed by growing relevant worm strains on HT115 bacteria transformed with a vector allowing for IPTG inducible expression of dsRNA against the *zyg-1* gene. Bacterial strains containing this RNAi vector were cultured overnight at 37°C, centrifuged, and the pellet was re-suspended in 1/10 of the original volume. 50uL of concentrated culture was spotted onto a nematode growth medium (NGM) plate with 1mM IPTG and 50ug/uL of kanamycin and the RNAi spot was allowed to grow overnight at 37**°**C. Plates were stored at 4°C for up to 10 days.

L4 hermaphrodite worms were transferred to RNAi plates, allowed to incubate for 2–3 hours, and then transferred to fresh RNAi plates. Live microscopy was performed on embryos 22–26 hours after worms were picked to the *zyg-1* RNAi plate. HT115 bacteria transformed with pHSG298 (Clontech) was used as a control for *zyg-1^RNAi^*.

### Auxin treatment

Auxin treatment was performed on worms 20–24 hours after L4 stage by transferring worms to bacteria-seeded plates containing auxin or ethanol. The natural auxin indole-3-acetic acid (IAA) was purchased from Alfa Aesar (#A10556). A 100 mM stock solution in ethanol was prepared and was stored at 4°C for up to two months. 100ul of the auxin stock solution or ethanol was added to NGM plates (volume 10mls) to achieve a final concentration of 1mM.

To perform RNAi against *zyg-1* and auxin treatment, L4 hermaphrodite worms were transferred to RNAi plates, allowed to incubate for 2–3 hours, and then transferred to fresh RNAi plates for 20-22 hours. Worms were then transferred to RNAi plates containing either auxin or ethanol to incubate for 2-3 hours before performing live microscopy.

## Statistical Analysis

Data was analyzed using Prism for statistical significance. For Figures 2C, D and E, student’s t-test was used to assess significance. For all other data, one-way ANOVA with Šídák correction was used to assess significance.

## Acknowledgements

We would like to thank Arshad Desai, Karen Oegema, Peter Askjaer and Geraldine Seydoux for valuable *C. elegans* strains. We thank the members of the Bhalla lab for reading of the manuscript. This work and the resulting manuscript was supported by the National Institutes of Health (NIH) (grant numbers R35GM141835 [N.B.], T32GM133391 [S.B.] and R01GM065591 [A.F.D.]), a postdoctoral fellowship of the Human Frontier Science Program (LT000903/2013-C [S.K.]) and the Howard Hughes Medical Institute (A.F.D.). Some strains were provided by the CGC, which is funded by NIH Office of Research Infrastructure Programs (P40 OD010440).

Because this manuscript is the result of funding in whole or in part by the NIH, this manuscript is subject to the NIH Public Access Policy. Through acceptance of this federal funding, NIH has been given a right to make this manuscript publicly available in PubMed Central upon the Official Date of Publication.

---

**Supplemental Figure 1:** A. Images of AB and P_1_ cells expressing mCherry::H2B (magenta) and GFP::PH (green) at NEBD or OCC in control or *zyg-1^RNAi^* embryos. B. Grayscale images of mCherry::H2B in control or *zyg-1^RNAi^* embryos. Scale bars indicate 5 microns.

**Supplemental Figure 2: Grayscale images of PCH-2::GFP in *AID::npp-21* strains without (top) and with TIR1 (bottom).** Area of enrichment indicated by yellow dotted circle. Scale bar indicates 5 microns.

**Supplemental Movie 1: In 75% of *C. elegans* two-cell embryos, NPP-21::GFP forms a spindle matrix only in P_1_ cells.** A two cell embryo expressing mCherry::H2B (red) and NPP-21::GFP (green) undergoing mitosis. The AB cell divides first and is on the right and the P_1_ cell divides after the AB cell and is on the left.

**Supplemental Movie 2: In 25% of *C. elegans* two-cell embryos, NPP-21::GFP forms a spindle matrix in both AB and P_1_ cells.** A two cell embryo expressing mCherry::H2B (red) and NPP-21::GFP (green) undergoing mitosis. The AB cell divides first and is on the right and the P_1_ cell divides after the AB cell and is on the left.

**Supplemental Table 1: *C. elegans* strains used in this study**

