## Supplemental Figure 1 for "NPP-21/TPR is required for developmental control of spindle checkpoint strength in *C. elegans*"

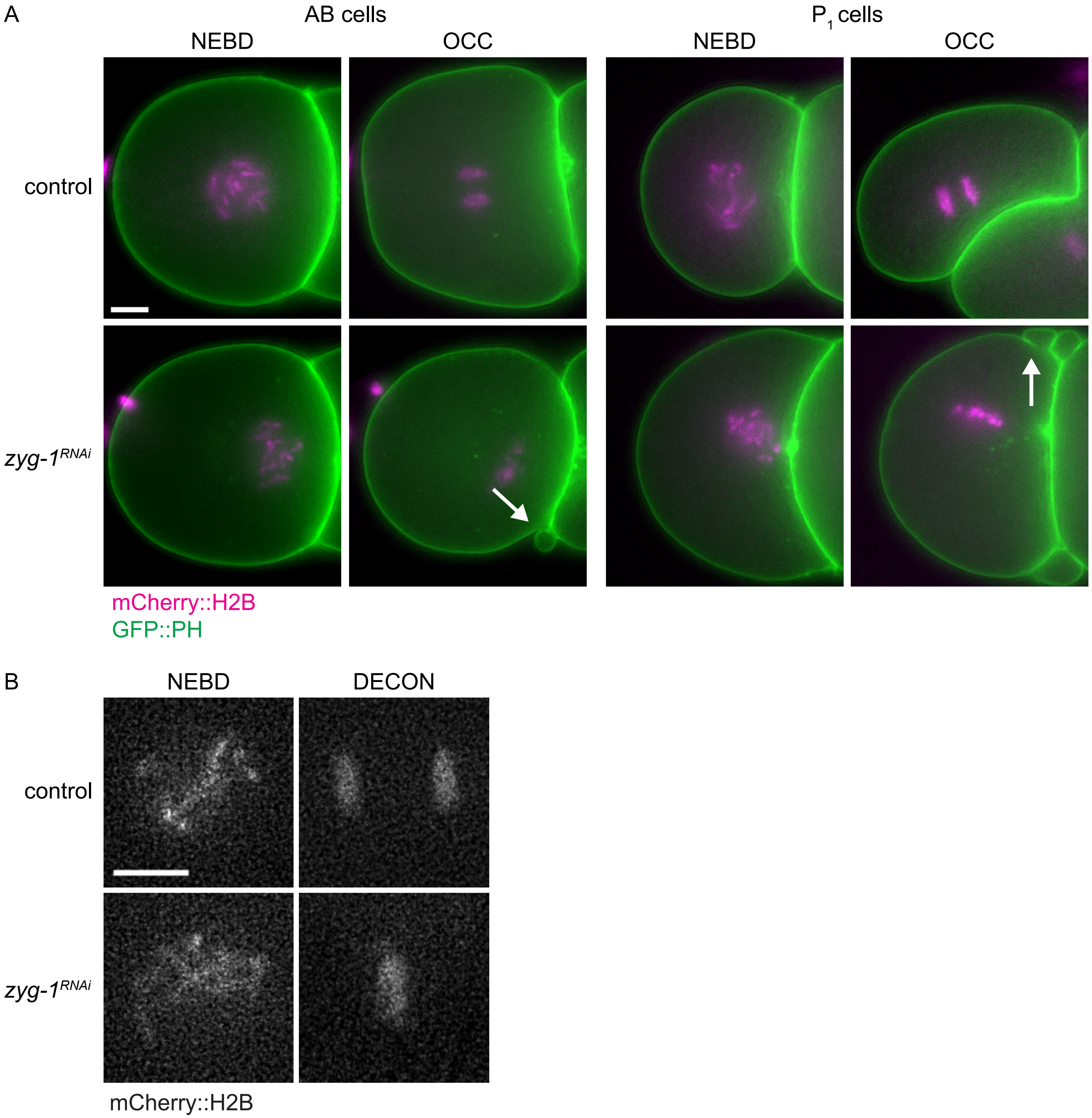


**Supplemental Figure 1: Hallmarks of mitotic entry and exit in *C. elegans* embryos.** A. Images of AB and $P_1$ cells expressing mCherry::H2B (magenta) and GFP::PH (green) at NEBD or OCC in control or *zyg-1*$^{RNAi}$ embryos. B. Grayscale images of mCherry::H2B at NEBD or DECON in control or *zyg-1*$^{RNAi}$ embryos. Scale bars indicate 5 microns.
