## Supplemental Figure 2 for "NPP-21/TPR is required for developmental control of spindle checkpoint strength in *C. elegans*"

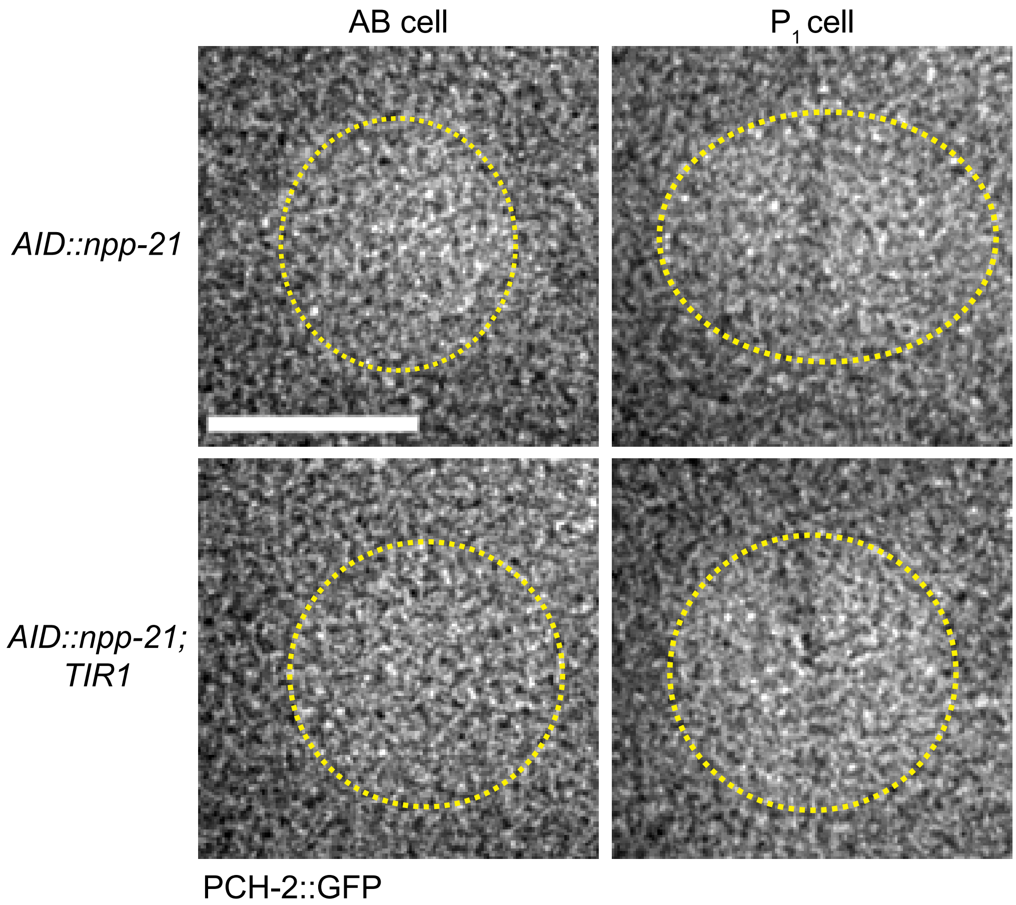


**Supplemental Figure 2: Grayscale images of PCH-2::GFP in *AID::npp-21* strains without (top) and with TIR1 (bottom).** Area of enrichment indicated by yellow dotted circle. Scale bar indicates 5 microns.
