## Supplemental Table 1 for "NPP-21/TPR is required for developmental control of spindle checkpoint strength in *C. elegans*"

**Supplemental Table 1. *C. elegans* strains used in this study**

| Strain Number | Genotype |
|---|---|
| OD595/BHL539 | *unc-119(ed3) III; ltIs37* [*pAA64; pie-1/mCherry::his-58; unc-119 (+)*] *IV; ltIs38* [*pAA1; pie-1/GFP::PH(PLC1delta1); unc-119(+)*] |
| BN1062/BHL114 | *npp-21::GFP(bq1) bqSi189[pBN13(unc-119(+) Plmn-1::mCherry::his-58)] II; unc-119(ed3) III* |
| CA1503/BHL1119 | *Pgld-1::TIR-1-mRuby(ieSi64) AID-3xFlag::npp-21(ie119) II* |
| BHL600 | *unc-119(ed3) III; ltIs37* [*pAA64; pie-1/mCherry::his-58; unc-119 (+)*] *IV; ltIs52* [*pOD379; pie-1/GFP::MDF-2; unc-119 (+)*] |
| BHL664 | *pch-2::GFP-3xFLAG(blt04, pCN94) II; unc-119(ed3) III; ltIs37* [*pAA64; pie-1/mCherry::his-58; unc-119 (+)*] *IV* |
| BHL1134 | *Pgld-1::TIR-1-mRuby(ieSi64) AID-3xFlag::npp-21(ie119) II; unc-119(ed3) III; ltIs37* [*pAA64; pie-1/mCherry::his-58; unc-119 (+)*] *IV; ltIs38* [*pAA1; pie-1/GFP::PH(PLC1delta1); unc-119(+)*] |
| BHL1177 | *Pgld-1::TIR-1-mRuby(ieSi64) AID-3xFlag::npp-21(ie119) II; unc-119(ed3) III; ltIs37* [*pAA64; pie-1/mCherry::his-58; unc-119 (+)*] *IV; ltIs52* [*pOD379; pie-1/GFP::MDF-2; unc-119 (+)*] |
| BHL1181 | *AID-3xFlag::npp-21(ie119) pch-2::GFP-3xFLAG(blt04, pCN94) II; unc-119(ed3) III; ltIs37* [*pAA64; pie-1/mCherry::his-58; unc-119 (+)*] *IV* |
| BHL1195 | *AID-3xFlag::npp-21(ie119) pch-2::GFP-3xFLAG(blt04, pCN94) II; unc-119(ed3) III; ltIs37* [*pAA64; pie-1/mCherry::his-58; unc-119 (+)*] *ieSi38 [Psun-1::TIR1::mRuby::sun-1 3'UTR, cb-unc-119(+)] IV* |
